# Sufficient therapeutic effect of cryopreserved frozen adipose-derived regenerative cells on burn wounds

**DOI:** 10.1101/436444

**Authors:** Yasuhiko Kaita, Takehiko Tarui, Hideaki Yoshino, Takeaki Matsuda, Yoshihiro Yamaguchi, Takatoshi Nakagawa, Michio Asahi, Masaaki Ii

**Author notes:** Corresponding author: Masaaki Ii, MD, PhD.

## Abstract

The purpose of this study was to evaluate whether cryopreserved (frozen) adipose-derived regenerative cells (ADRCs) have a therapeutic effect on burn wound healing as well as freshly isolated (fresh) ADRCs.

Full thickness burns were created on dorsum of nude mice and burn wound was excised. The wound was covered by artificial dermis with; (i) fresh ADRCs, (ii) frozen ADRCs, and (iii) PBS (control). The assessment for wound healing was performed by morphological, histopathological and immunohistochemical analyses.

In vivo analyses exhibited the significant therapeutic effect of frozen ADRCs on burn wound healing up to the similar or higher level of fresh ADRCs. There were significant differences of wound closure, epithelized tissue thickness, and neovascularization between the treatment groups and control group. Although there was no significant difference of therapeutic efficacy between fresh ADRC group and frozen ADRC group, frozen ADRCs improved burn wound healing process in dermal regeneration with increased great type I collagen synthesis compared with fresh ADRCs.

These findings indicate that frozen ADRCs allow us to apply not only quickly but also for multiple times, and the cryopreserved ADRCs could therefore be useful for the treatment of burn wounds in clinical settings.

## Introduction

Severe burns can be fatal despite recent advances in burn care [1]. For saving life in severe burn patients, early wound closure is one of the most important factors. Thus, the improvement of burn wound treatment is still required. Many types of stem cells have been studied for use in a variety of diseases including burns [2–5]. Over the last decade, the application of mesenchymal stem cells (MSCs) has been emerged as a promising therapy for wound healing. The MSCs, which can be isolated from bone marrow, adipose tissue, umbilical cord/blood, and amnion, exhibit favorable effects on wounds and play a central role in healing process [6–8]. Among these MSCs, adipose tissue-derived MSCs exhibit the advantages of harvesting source tissue with less invasive procedure and easy culture expansion, specifically, in case of patients with burns who need to undergo a certain surgical treatment [9].

Adipose-derived stem cells (AdSCs) have recently been shown to accelerate wound healing in various animal models [10–12]. In particular, AdSCs decreased the wound surface area in acute burns in several studies. AdSCs increased fibroplasia and collagen remodeling in the process of wound healing [13]. AdSCs also enhanced vascularity, collagen deposition, and adipogenesis in burn wound healing[14]. The stromal vascular fraction (SVF) isolated from enzymatically digested adipose tissue, which is called adipose-derived regenerative cells (ADRCs), was reported to decrease inflammation and increase vascularization in rats with acute burns [15]. The treatment for burn wounds with uncultured adipose-derived stromal cells which are equivalent to ADRCs in combination with artificial dermis enhanced angiogenesis and induced blood vessel maturation resulting in matrix remodeling in the regenerated tissue [16]. These studies revealed the favorable effect by a single administration/transplantation of freshly isolated SVF cells (ADRCs) on acute burns. Nevertheless, cryopreservation of the freshly isolated ADRCs is required for multiple administrations/transplantations to obtain better outcomes. Recently, stem cell banking system has been developed for future autologous or allogeneic stem cell therapy for diseases including burns, however, the potential efficacy of the cryopreserved ADRCs after thawing have not been investigated sufficiently [17].

We therefore examined the therapeutic effects of cryopreserved ADRCs, which can be isolated from aspirated or surgically resected fat tissue and stored in large numbers, on burn wounds, and tested our hypothesis that the cryopreserved ADRCs displayed advantages of multiple regenerative cell therapy. In the present study, we examined the functions of ADRCs and compared the effects of freshly isolated ADRCs with the cryopreserved frozen ADRCs on burn wounds evaluating the skin tissue regeneration following debridement.

## Materials and Methods

### Murine burn wound model

Six- to eight-week-old male BALB/c nude mice (SHIMIZU Laboratory Supplies, Kyoto, Japan) were used for the experiments. The animals were housed in individual cages, and wounds were created using previously described methods [13]. Briefly, the mice were anesthetized with an intraperitoneal injection of Avertin™ (200 mg/kg, Ben Venue Laboratories, Bedford, OH, USA). Burn wounds were created by placing a pre-heated (150°C) aluminum column (custom made; 6 mm in diameter) on the dorsum for 5 seconds after shaving hair. The heat-induced injuries were verified as a full thickness burn by histological findings. All animals were intraperitoneally administered 0.5 ml of Lactated Ringer’s Solution (Lactec™, Fuso, Osaka, Japan) to avoid acute dehydration following the stress of burn procedure.

### Surgical procedures

The advanced burn causes tissue necrosis and the burn tissue is generally excised as a therapeutic approach for patients. We therefore excised the heat-rod contacted skin tissue mimicking clinical settings. One hour after burn creation, mice were anesthetized and burn wounds were excised. After the damaged skin had been excised, the edges of the wound were sutured to the fascia to prevent skin shrinking. The wound was covered with an artificial dermis (Terudermis™, Olympus Terumo Biomaterials Corp., Tokyo, Japan) and (i) fresh ADRCs (5 × 10^4^ cells/30 μl of PBS; fsADRC group, n = 6), (ii) frozen ADRCs (5 × 10^4^ cells/30 μl of PBS; fzADRC group, n = 6), or (iii) PBS (30 μl; control group, n = 6). A semipermeable transparent dressing (Tegaderm; 3M Health Care, St. Paul, MN, USA) was placed over the wound and sealed at the edges with Compound Benzoin Tincture (3M™ Steri-Strip™ Compound Benzoin Tincture C1544; 3M Health Care, St. Paul, MN, USA) to prevent the wound contracture in addition to the suture fixation described above. All animals were intraperitoneally administered 0.5 ml of Lactated Ringer’s Solution following the surgical procedure. ADRCs were directly loaded on the artificial dermis at a density of 1.8 × 10^5^ cells/cm^2^. For this purpose, Terudermis™ was cut into a 6-mm diameter circles and placed on the dish with the silicone layer facing down. Next, 30 μl of PBS containing 5 × 10^4^ ADRCs were soaked on the surface of Terudermis, and then the ADRC-loaded artificial dermis was placed in direct contact with the wound. At day 6 and 12, wounds were harvested and analyzed by Western blotting and quantitative real-time reverse transcription polymerase chain reaction (qRT-PCR) to assess wound healing.

### ADRC isolation and cryopreservation

Human subcutaneous adipose tissues were obtained from healthy female donors undergoing elective liposuction with informed consent. Adipose tissues were minced and digested with Celase (1 U/ml; Cytori Therapeutics, San Diego, CA, USA) for 20 min at 37°C. After filtration through a 100-μm cell strainer (BD Biosciences, San Jose, CA, USA), ADRCs were isolated by centrifugation (300 *g* for 5 min) removing adipocytes. Total cell number and cell viability were measured with a LUNA automated cell counter (Logos Biosystems, Inc., USA). The freshly isolated ADRCs were maintained in Lactated Ringer’s Solution for in vitro experiments, including assessments of gene expression, and were suspended in STEM-CELLBANKER (Nippon Zenyaku Kogyo Co., Fukushima, Japan) for cryopreservation at −80°C. The frozen ADRCs were thawed, followed by washing with PBS, and incubated in Lactated Ringer’s Solution for 6-h recovery period at 37°C and used in the assessment of gene expression. The freshly isolated cells (fresh) ADRCs from fat tissue and the cryopreserved (frozen) ADRCs were used for all experiments without passage.

### Characterization of fresh ADRCs and frozen ADRCs

The cells (5 × 10^5^) were incubated with fluorescent dye-conjugated mouse monoclonal antibodies against CD90, CD105, CD29, CD34, CD14, and CD45 (1:1000; BioLegend, San Diego, CA, USA) for 30 minutes at 4°C. Flow cytometry was performed to characterize the ADRC phenotypes using a Cell Analyzer EC800 (SONY, Tokyo, Japan), according to the manufacturer’s instructions.

### Cell functional assays with ADRC-derived conditioned medium

The migration and proliferation of keratinocytes and fibroblasts were examined following treatment with ADRC-derived conditioned medium (CM) to evaluate the paracrine effects of fresh and frozen ADRCs on wound healing. Freshly isolated ADRCs and thawed frozen ADRCs were plated on 100-mm dishes and incubated with 1% FBS/DMEM-F12 medium to prepare ADRC-derived CM. After 48 h in culture, the medium was collected for experiments. Normal human epidermal keratinocytes (NHEKs) and normal human dermal fibroblasts (NHDFs) were purchased from KOHJIN BIO, Inc. (Saitama, Japan) and Takara Biochemicals, Inc. (Kyoto, Japan), respectively, and were cultured in 10% FBS/DMEM-F12 medium (FBS; Gibco; Thermo Fisher Scientific, Inc., Grand Island, NY, USA).

A modified Boyden’s chamber was used for migration assay, as previously described [18]. Briefly, a polycarbonate filter (5-μm pore size) (Transwell™) was placed between the upper and lower chambers. NHEK and NHDF suspensions (5 × 104 cells/well) were placed in the upper chamber, and the lower chamber was filled with 0% FBS/DMEM-F12, 20% FBS/DMEM-F12, fresh ADRC-derived CM, and frozen ADRC-derived CM. The cells were incubated for 6 h at 37°C in a 5% CO_2_ incubator. Migration was evaluated by calculating the number of cells that migrated through the polycarbonate filter to the lower chamber in each well. The number of migrated cell was counted in 5 randomly selected high-power fields and averaged.

Cell Counting Kit-8 (DOJINDO, Tokyo, Japan) was used for the proliferation assay. Briefly, NHEKs and NHDFs (5000 cells/well) were seeded on 96-well flat-bottomed plates with 100 μl of growth medium: 1% FBS/DMEM-F12, 10% FBS/DMEM-F12, fresh ADRC-derived CM, or frozen ADRC-derived CM. Then, the cells were incubated for 48 h at 37°C in a 5% CO_2_ incubator. The absorbance was recorded at 570 nm using a 96-well ELISA plate reader (SPECTRA MAX 190, Japan Molecular Device). All experiments were performed in triplicate and confirmed the reproducibility.

### Quantitative real-time RT–PCR analysis

The mRNA expression levels of the epidermal growth factor (*EGF*), fibroblast growth factor 2 (*FGF2*), hepatocyte growth factor (*HGF*), and vascular endothelial growth factor (*VEGF*) were examined by quantitative real-time reverse transcription polymerase chain reaction (qRT-PCR) to confirm the effects of ADRCs on dermal regeneration. Fresh and frozen ADRCs (5 × 10^5^ cells/dish) were plated on 35-mm culture dishes and incubated for 48 h at 37°C in a 5% CO_2_ incubator. In addition, mRNA levels of the type I collagen and type III collagen were analyzed with tissue sample by qRT-PCR for the assessment of dermal regeneration. Total RNA was extracted from ADRCs using a RNeasy Mini Kit (QIAGEN Science, Hilden, Germany) and reverse transcribed using a ReverTra Ace™ qPCR RT Master Mix (TOYOBO Life Science, Osaka, Japan), according to the manufacturers’ instructions. For qRT-PCR, the converted cDNA samples (2 μl) were amplified in triplicate with a real-time PCR machine (CFX Connect™, Bio-Rad, Hercules, CA, USA) in a final volume of 10 μl using SYBR Green Master Mix reagent (Bio-Rad) and gene-specific primers for human samples (Table 1) and mouse samples (Table 2). A melting curve analysis was performed with Dissociation Curves software (Bio-Rad), and the mean cycle threshold values were used to calculate gene expression levels that were normalized to human glyceraldehyde 3-phosphate dehydrogenase (*GAPDH*) mRNA levels.

**Table 1.**
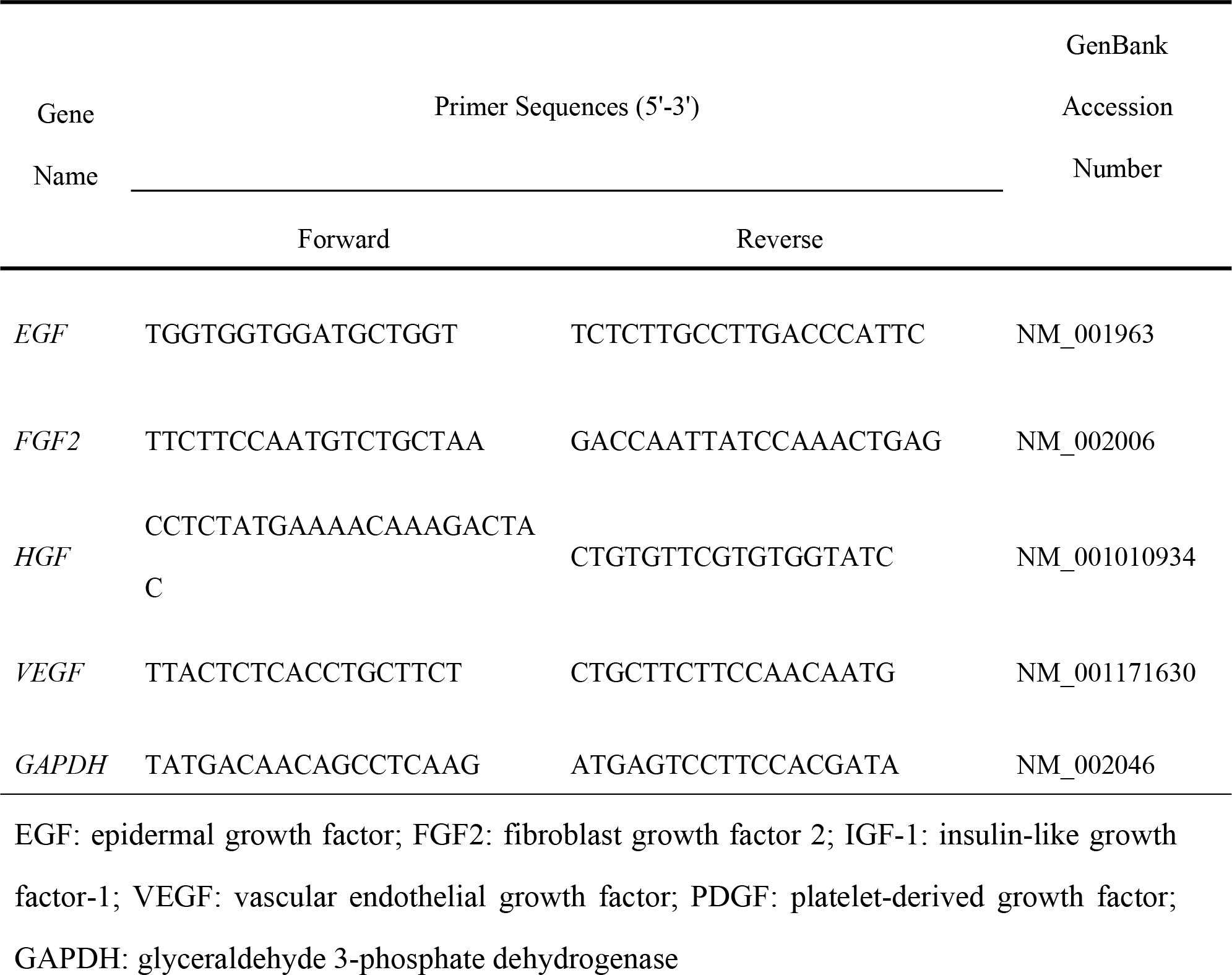
Specific primers for human samples used for qRT-PCR.

**Table 2.**
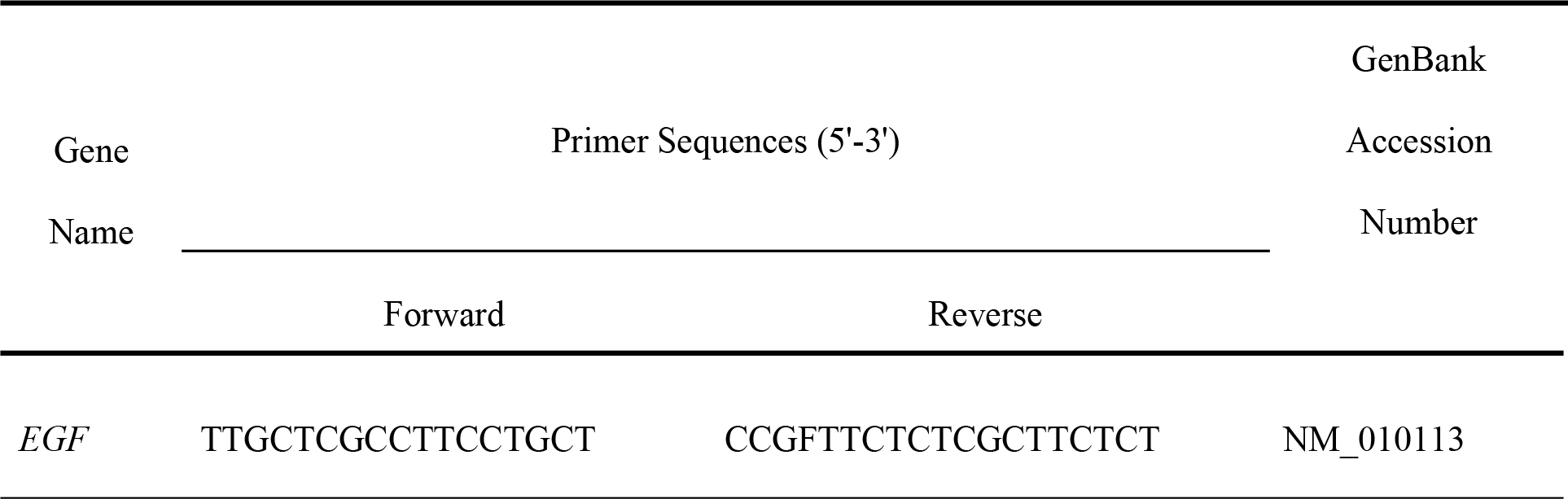
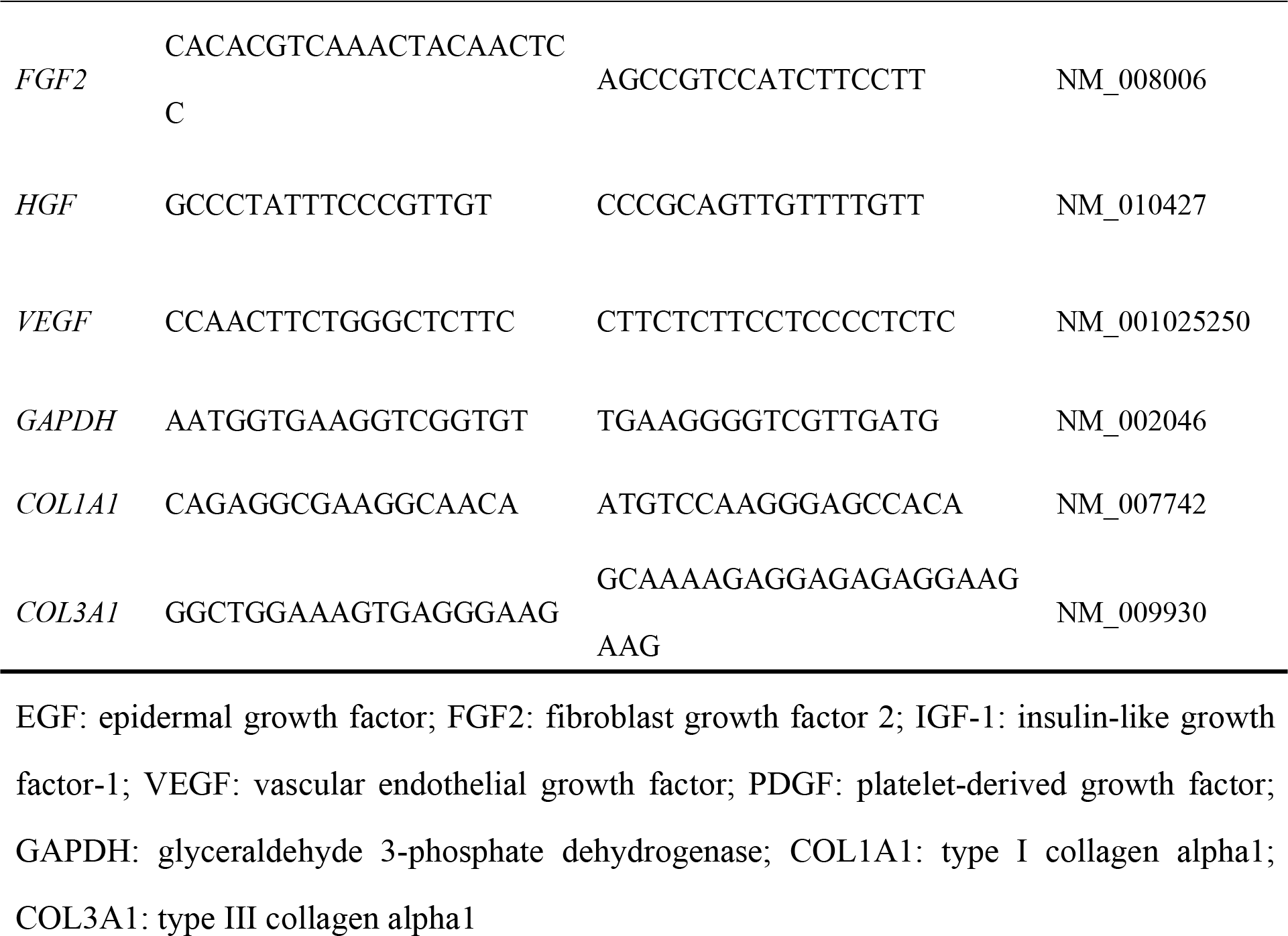
Specific primers for mouse samples used for qRT-PCR.

### Wound closure measurements

The treated mice were observed, and digital images of the dorsal side were captured on days 0, 6, and 12. The wound area was measured by tracing the margins and was expressed as pixel area using image analysis software (ImageJ, NIH, USA). The measurements were performed in triplicate and were evaluated as the percentage of closure from the original wound using the following formula: % of wound closure = 100 × (wound area on day 0 – wound area on day 6 and 12) / wound area on day 0.

### Histological analysis

At day 12, wounds were harvested and fixed with 4% neutral-buffered paraformaldehyde (PFA) for normal histopathological analysis. Frozen tissue sections (6 μm) embedded in optimum cutting temperature (O.C.T.) compound (Tissue-Tek™, Sakura, Japan) were stained by Masson’s trichrome method. The thickness of the regenerated skin and original skin thickness were measured to evaluate dermal regeneration. The skin thickness was measured at three points in each site and averaged. The ratio of the thickness of the regenerated skin to the original skin thickness was calculated to evaluate tissue regeneration using an image analysis software (ImageJ, NIH, USA). To evaluate collagen production in the wounds, the skin tissue samples at day 6 and 12 were analyzed by Picro-Sirus Red staining. The tissue sections were incubated in Picro-Sirus Red Solution (Scy Tek Laboratories, Inc. Logan, UT) for 1hour at room temperature. After washing in acetic acid solution for two times, the slides were dehydrated through absolute ethanol and mounted in synthetic resin. The sections were examined under a standard light and polarized light microscopy. The area of collagen fibers was calculated for the evaluation of collagen synthesis using an image analysis software (Image J™, NIH special volunteer, USA).

### Immunohistochemistry

The wounded skin tissues were harvested at day 6 and 12 and fixed with 4% PFA in PBS overnight at 4°C and quenched in 20% sucrose/PBS overnight at 4°C. The tissues were embedded in O.C.T. compound (Tissue-Tek™, Sakura, Japan) and frozen (−80°C) for cryosectioning. For assessment of neovascularization, the tissue sections were incubated with a biotinylated primary antibody against isolectin B4 (ILB4, 1:100) (Vector Laboratories, Burlingame, CA, USA) diluted in PBS containing calcium and magnesium for 1 h at room temperature, followed by 15 min-incubation with a DyLight 549-conjugated streptavidin secondary antibody (1:1000; Vector Laboratories, Burlingame, CA, USA). In order to assess the retention of the transplanted ADRCs, the tissue sections were incubated with an anti-human mitochondria antibody (1:500, Abcam, USA) for 1 h at room temperature, followed by 15 min-incubation with a DyLight 549-conjugated secondary antibody (1:1000; Vector Laboratories, Burlingame, CA, USA). The tissue sections serving as controls were stained without the primary antibody. The nuclei were counterstained with DAPI. The tissue sections were examined under a fluorescent microscope (BZ-8000, Keyence, Osaka, Japan). Capillaries and large vessels were counted in a high-power field (X100), and the vascular density was calculated and averaged in each experimental group to evaluate neovascularization.

### Western blot analysis

The regenerated skin tissue was harvested at day 6 and day 12 and lysed in sodium dodecyl sulfate-polyacrylamide gel electrophoresis (SDS-PAGE) sample buffer. The proteins in lysates (5 μg) were loaded for each condition and resolved using SDS-PAGE. Separated proteins were electrophoretically transferred to poly vinylidene di-fluoride (PVDF) membranes (Merck Millipore, Darmstadt, Germany), blocked with 5% skim milk, and probed with an antibody for type III collagen (1:1000, Bioworld Technology, Inc. Loius Park, MN, USA) and ◻-tublin (1:1000, Sigma-Aldrich Japan, Tokyo, Japan). The appropriate horseradish peroxidase-conjugated secondary antibodies were used (Jackson Laboratory, Bar Harbor, ME, USA). Signals were detected and documented using an LAS3000 densitometry system (Fujifilm, Tokyo, Japan) or a Fusion FX7 (Vilber Lourmat, Eberhardzell, Germany).

### Ethical considerations

This study was conducted according to the guidelines and with the approval of the ethics committees of the Osaka Medical College (reference number: 1040-01) and Sobajima Clinic. All patients consented to the use of their tissues. The Institutional Animal Care and Use Committee of Osaka Medical College approved all research protocols (approval ID: 28081), including the surgical procedures and animal care procedures.

### Statistical analysis

All values are presented as means ± standard errors of the means (SEM). Statistical analyses were performed with commercially available software (GraphPad Prism™, MDF Co. Ltd., Japan). Comparisons between 2 groups were performed using Student’s t-test, and comparisons among multiple groups were assessed for significance using analysis of variance (two-way ANOVA), followed by post-hoc testing with Tukey’s test. P values less than 0.05 were considered statistically significant.

## Results

### Frozen ADRCs expressed high levels of MSC markers

The surface markers of ADRCs obtained from 4 donors were analyzed to characterize the cell populations. Based on the results by flowcytometry, frozen ADRCs comprised greater proportions of CD90- and CD29-positive cells than fresh ADRCs, suggesting that cells other than MSCs perished during cryopreservation and thawing. On the other hand, there was no significant difference in Leukocytes (CD45+), monocytes (CD14+), and hematopoietic stem cells (CD34+) between the fresh and frozen ADRC groups. Thus, frozen ADRCs comprised a heterogeneous cell population, including stem cells and leukocytes, similar to fresh ADRCs. (Figure 1)

**Figure 1.**
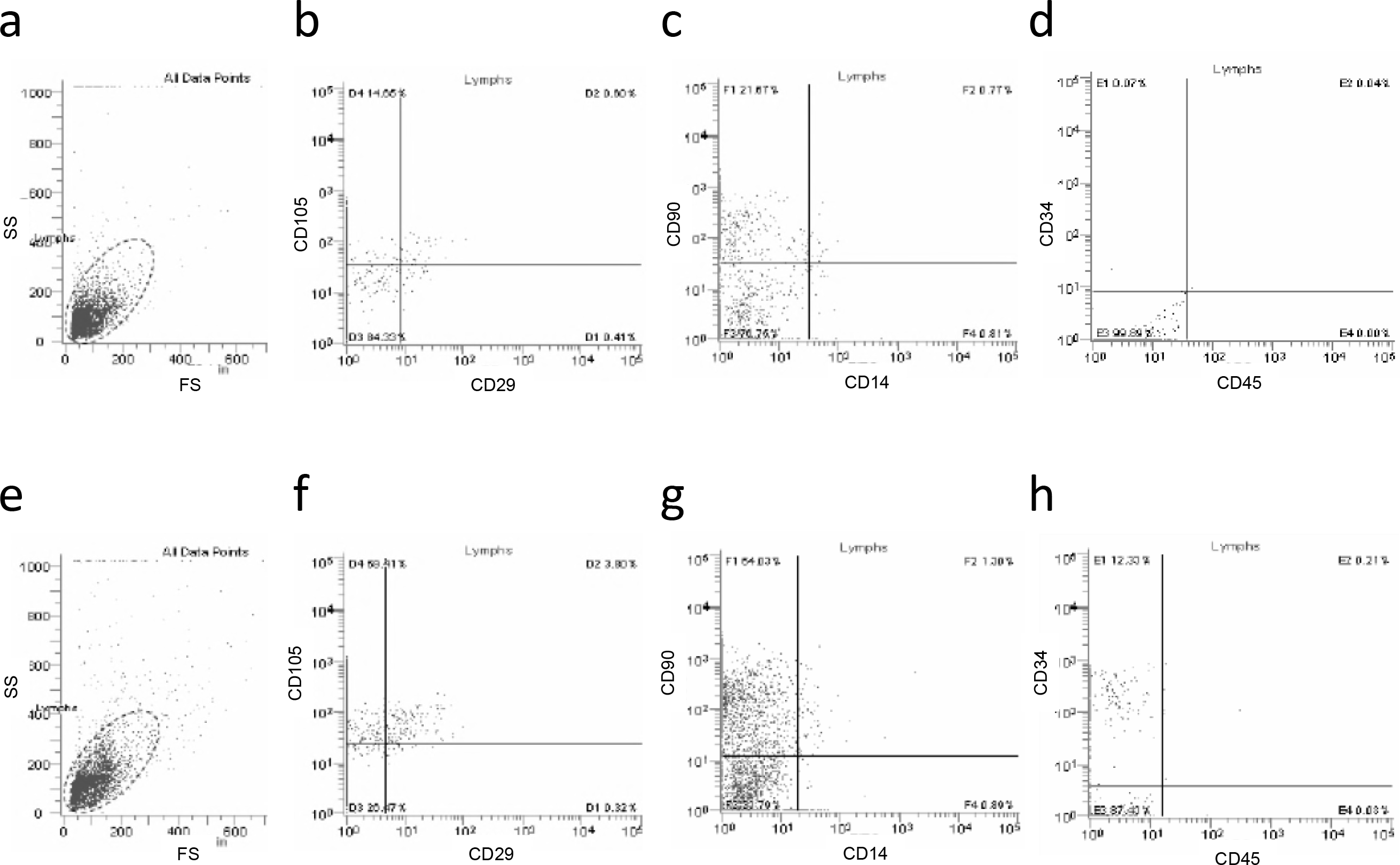
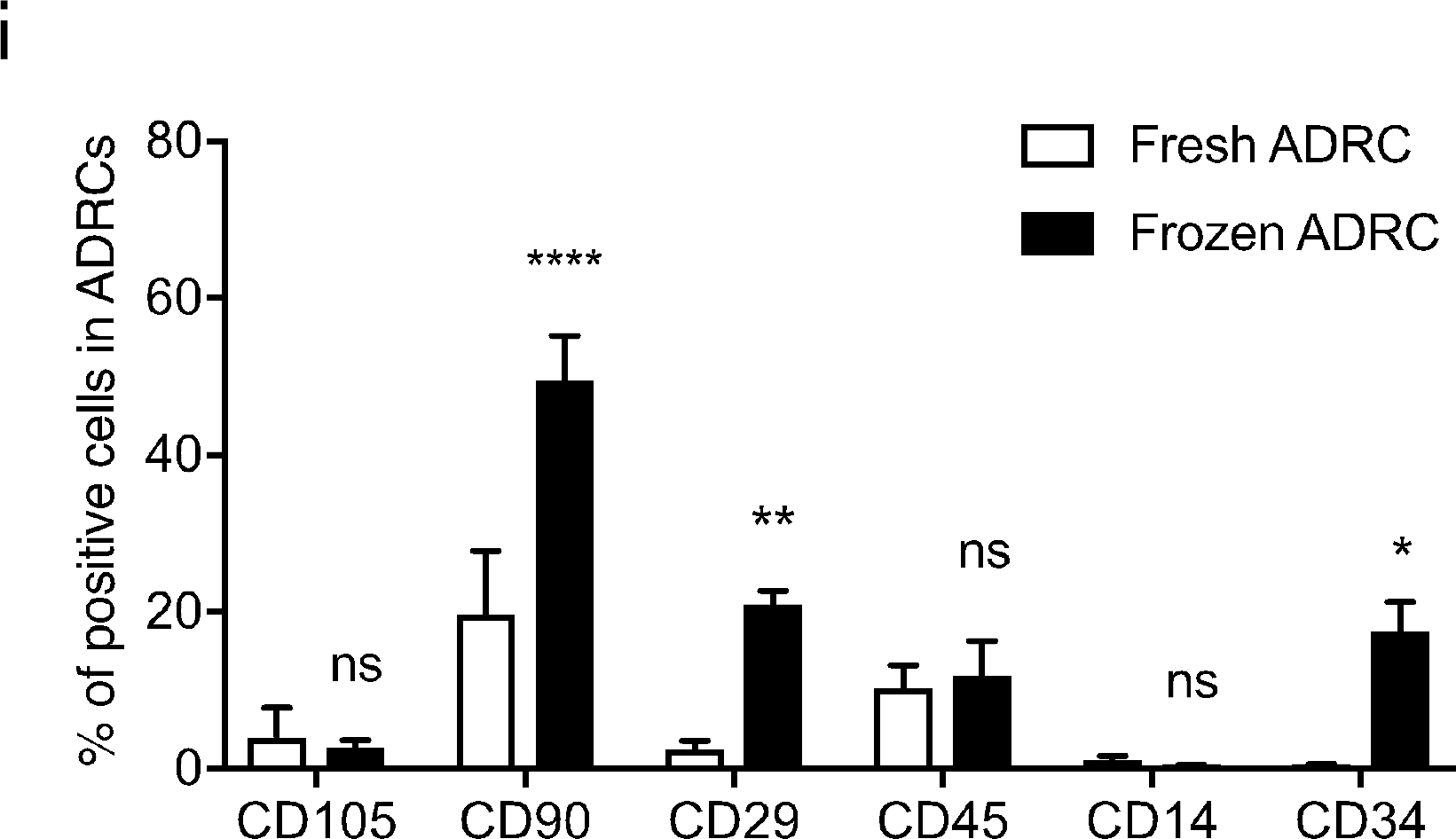
Assessment of the characteristics of fresh and frozen ADRCs. A and E, Fresh and frozen ADRCs were analyzed based on forward scatter (FS) and side scatter (SS). B~D, and F~H, FACS analysis of CD105, CD29, CD90, CD19, CD34, and CD45 expressions in fresh and frozen ADRCs. I, Flow cytometry analysis of the expression levels of CD markers in fresh and frozen ADRCs. The percentages of CD90-, CD105-, CD29-, CD34-, CD14-, and CD45-positive cells in fresh and frozen ADRCs are presented in the bar graph. *, p < 0.05; **, p < 0.01; and ****, p < 0.0001 compared with fresh ADRCs.

### Frozen ADRC-derived CM stimulated proliferation and migration activities of keratinocytes and fibroblasts

NHEKs and NHDFs were cultured with ADRC-derived CM to examine whether ADRC-derived CM stimulated skin cell proliferation and migration during wound closure. A significant difference in the proliferation of NHEKs was observed among the 20% FBS group, frozen ADRC-derived CM group, and control group. A marked increase in the number of NHEKs was observed following treatment with fresh ADRC-derived CM compared to control treatment, but the difference was not significant. There were significant differences in the proliferation of NHDFs among 20% FBS group, fresh ADRC-derived CM group, frozen ADRC-derived CM group, and control group. There was a significant difference of proliferation in NHEKs and NHDFs between fresh ADRC-derived CM group and frozen ADRC-derived CM group. (Figure 2A and 2B)

**Figure 2.**
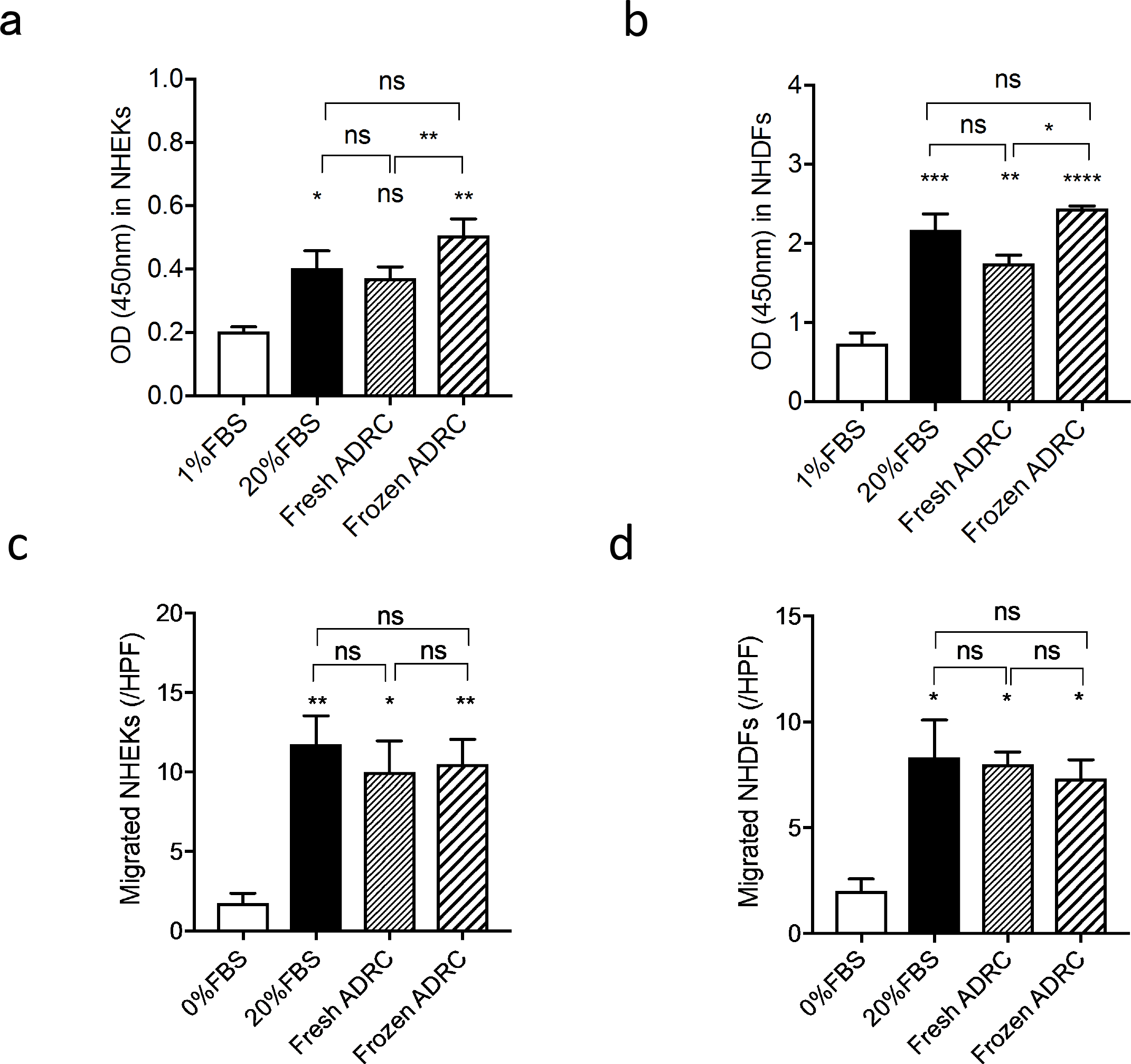
Assessment of the functions of fresh and frozen ADRCs. A and B, Proliferation of keratinocytes and fibroblasts co-cultured with fresh ADRCs or frozen ADRCs. The proliferation of NHEKs (A) and NHDFs (B) is expressed as the optical density (OD) at 450 nm. The cells cultured in 1% FBS/DMEM-F12 and 20% FBS/DMEM-F12 were evaluated as the negative and positive controls, respectively. ns, not significant; *, p < 0.05; **, p < 0.01; ***, p < 0.001; and ****, p < 0.0001 compared with 0.5% FBS. C and D, Migration of NHEKs and NHDFs co-cultured with fresh ADRCs or frozen ADRCs. The migration of NHEKs (C) and NHDFs (D) is presented as the numbers of migrated cells in a high-power field (/HPF). ns; *, p < 0.05; and **, p < 0.01 compared with 1% FBS.

A significant difference in the migration of NHEKs was observed among 20% FBS group, fresh ADRC-derived CM group, frozen ADRC-derived CM group, and control group. However, significant difference was not observed between fresh ADRC-derived CM group and frozen ADRC-derived CM group. There were significant differences in the migration of NHDFs among 20% FBS group, fresh ADRC-derived CM group, frozen ADRC-derived CM group, and control group. However, there was no significant difference between the fresh ADRC-derived CM group and frozen ADRC-derived CM group. (Figure 2C and 2D) Based on these results, ADRCs appears to have sufficient potential to stimulate proliferation and migration activities of NHEKs and NHDFs those play an important role for wound healing.

### Frozen ADRCs expressed high levels of growth factor genes

The expression levels of growth factor genes in fresh ADRCs and frozen ADRCs were examined by qRT-PCR using specific primers for human *EGF*, *FGF2*, *HGF*, and *VEGF* to explore the paracrine mechanisms of ADRCs in skin regeneration. Both fresh and frozen ADRCs expressed high levels of the *FGF2* and *VEGF* mRNAs (Figure 3B and 3D).

**Figure 3.**
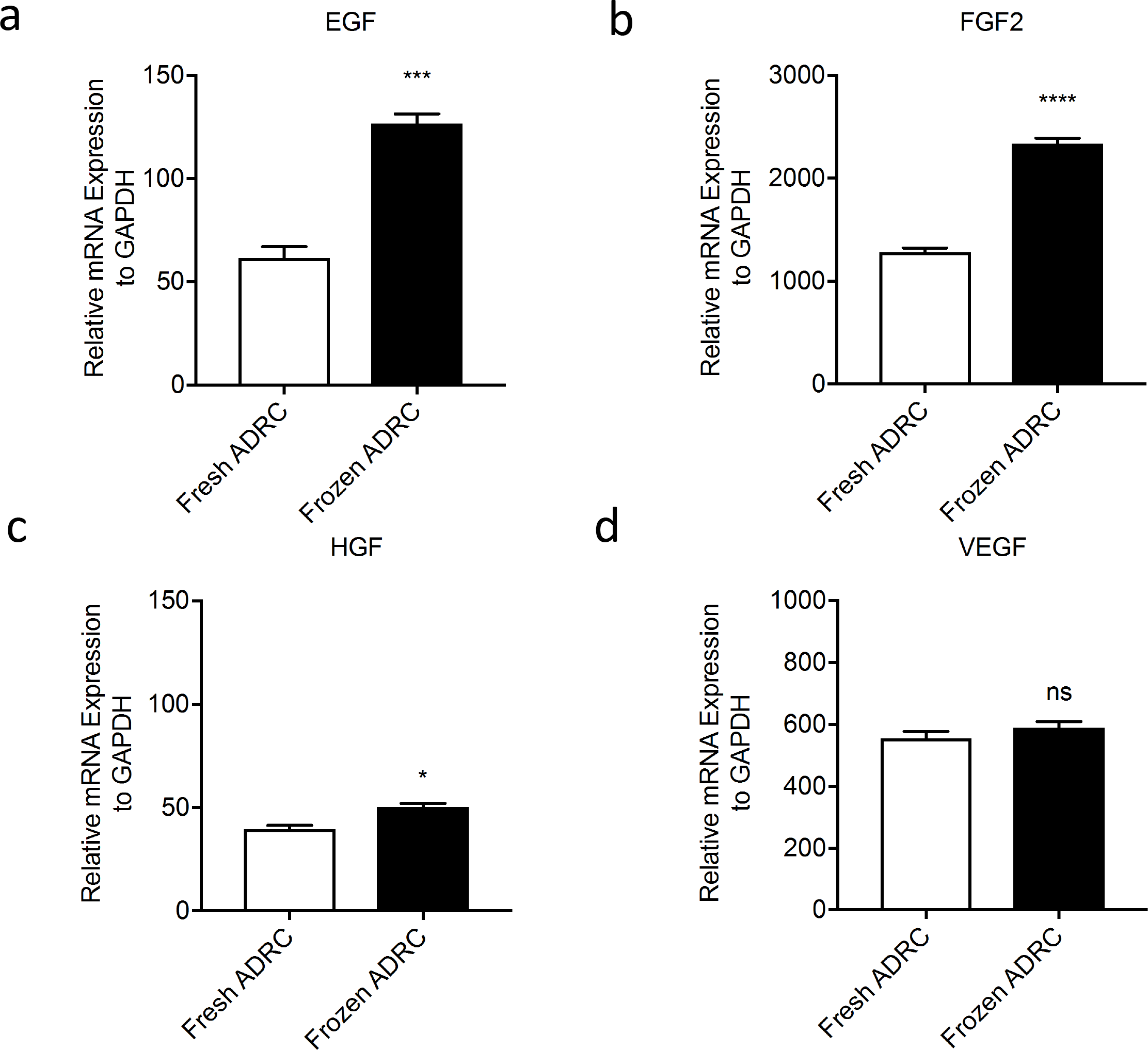
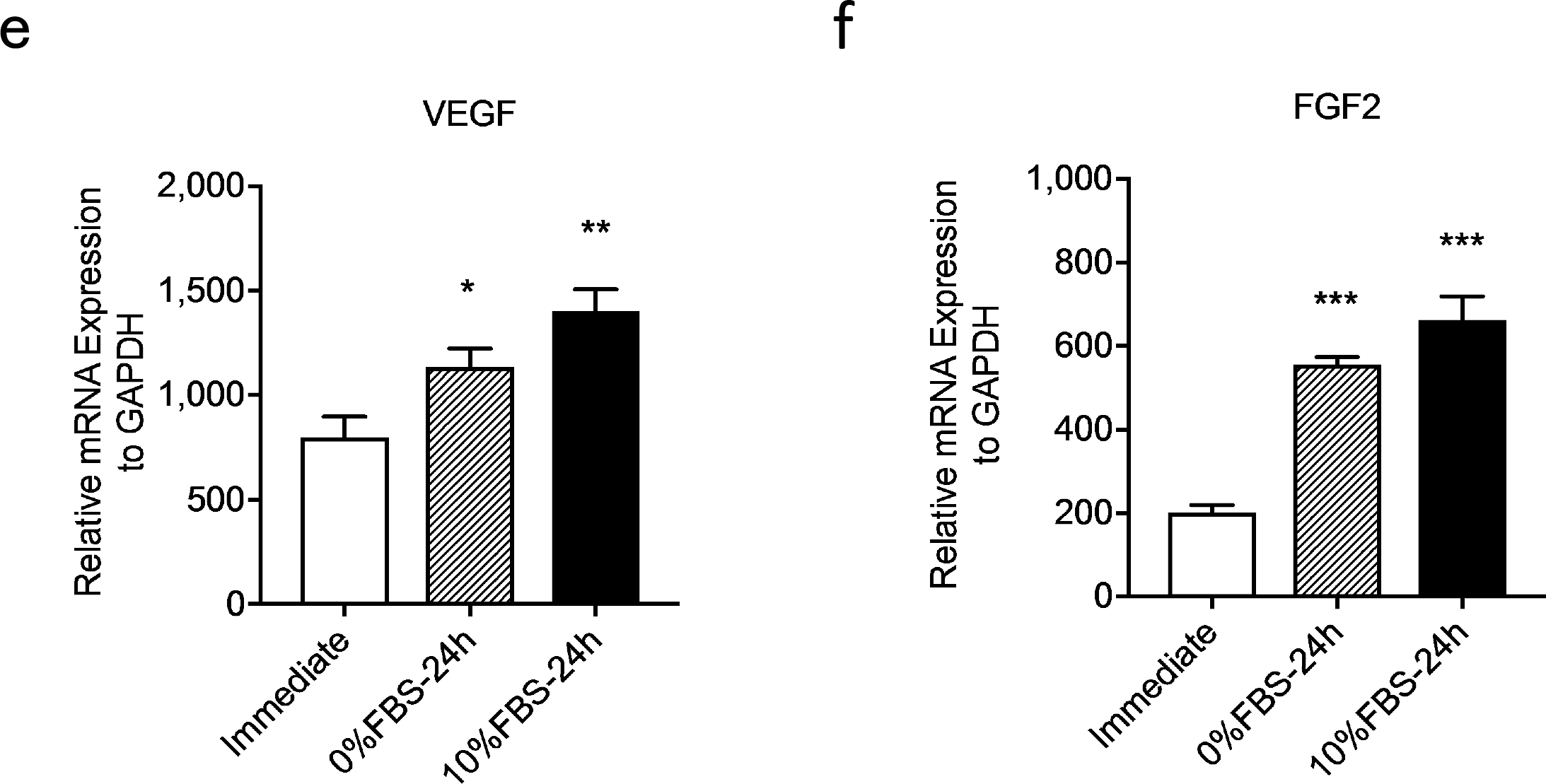
Expression of tissue regeneration-related growth factor genes in ADRCs. The mRNA expression levels in fresh ADRCs and frozen ADRCs, and frozen (cryopreserved) ADRCs immediately after thaw, those cultured with 0%FBS/DMEM F12 for 24h, and those cultured with 10%FBS/DMEM F12 for 24h were examined by quantitative real-time RT-PCR analysis using specific primers for the human *EGF* (A), *FGF2* (B and F), *HGF* (C), and *VEGF* (D and E) genes. The relative mRNA expression levels of target genes were normalized to the levels of the *GAPDH* mRNA. ns; *, p < 0.05; **, p<0.01; ***, p < 0.001; and ****, p < 0.0001 compared with the fresh ADRC group.

Although the *EGF* mRNA was not expressed at high levels (Figure 3A) in fresh ADRCs and frozen ADRCs, a significant difference was observed between fresh ADRCs and frozen ADRCs. The *EGF* and *HGF* mRNAs were expressed at significantly higher levels (Figure 3A and 3C) in frozen ADRCs than in fresh ADRCs; however, the *HGF* mRNA was expressed at low levels in fresh ADRCs and frozen ADRCs, and the difference in the expression was not significant. The expression of the *VEGF* mRNA (Figure 3D) was not significantly different between fresh ADRCs and frozen ADRCs. Thus, ADRCs promoted wound healing by secreting growth factors, and the paracrine effects of frozen ADRCs might be more effective than those of fresh ADRCs.

Next, we examined the gene expressions among cryopreserved ADRCs immediately after thaw, those cultured in 0%FBS medium for 24h, and those cultured in 10%FBS medium for 24h. The mRNA expressions of *VEGF* and *FGF2* were increased after 24h culture regardless of FBS supplementation in culture medium, although those of *EGF* and *HGF* were not detected in all groups. (Figure 3E and 3F) Based on these results, cryopreserved ADRCs cultured for 24h but not those immediately after thaw would exhibit more therapeutic efficacy, however, the process of cell culture may reduce the advantage of quick/simple application of cryopreserved cells in terms of clinical use, specifically, in case of emergency.

### Frozen ADRCs reduced burn wound surface area

All treatments were well tolerated by the animals (n = 6 in each group), and no evident side effects were observed in the control and treatment groups. The planimetry of the wounds was evaluated to investigate the therapeutic effects of ADRCs on burn wounds using representative digital images of wound healing progression at 0, 6, and 12 days post-surgery (Figure 4A). The percentage of wound closure was also analyzed (Figure 4B). No differences were observed among the different treatments on day 6. At day 12, a significant difference in the percentage of wound closure was observed between the treatment groups and PBS group. However, no significant differences were observed between fresh ADRC group and frozen ADRC groups. Based on these findings, ADRCs promoted burn wound healing, and frozen ADRCs exerted similar therapeutic effects on burn wound healing as fresh ADRCs.

**Figure 4.**
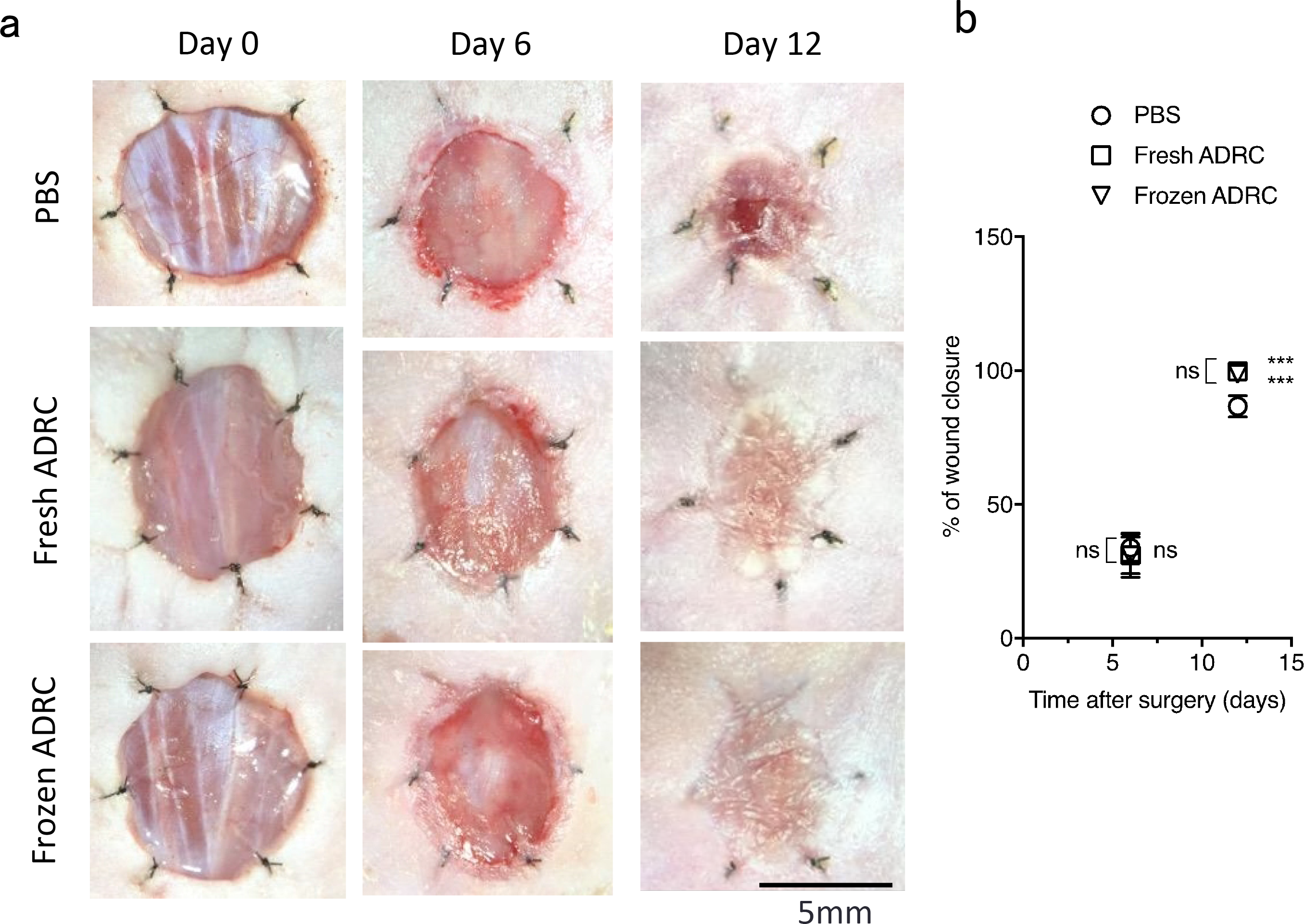
Effects of ADRCs on burn wound healing. A, Digital images of wounds during the healing process. The images are shown at 0, 6, and 12 days after surgery and treatments. B, Analysis of wound closure. The percentage of wound closure was calculated on days 6 and 12. (100 × (wound area on day 0 − wound area on day n) / wound area on day 0). ns; *, p < 0.05; **, p < 0.01; and ***, p < 0.001 compared with the PBS group (control).

### Frozen ADRCs promoted skin regeneration

Masson trichrome staining was used to evaluate skin regeneration morphometrically. The thickness of the regenerated tissue was compared with the thickness of the original skin. The representative images show epithelized and regenerated skin tissue and original skin tissue (Figure 5A). The ratio of the thickness of the regenerated skin tissue to the thickness of the original skin tissue was calculated. A significant difference in tissue thickness was observed between the treatment groups and the PBS group. However, no significant differences were observed between fresh ADRC group and frozen ADRC group. (Figure 5B) Thus, ADRCs induced skin regeneration, and frozen ADRCs improved skin tissue regeneration in burn wounds up to the similar level to fresh ADRCs.

**Figure 5.**
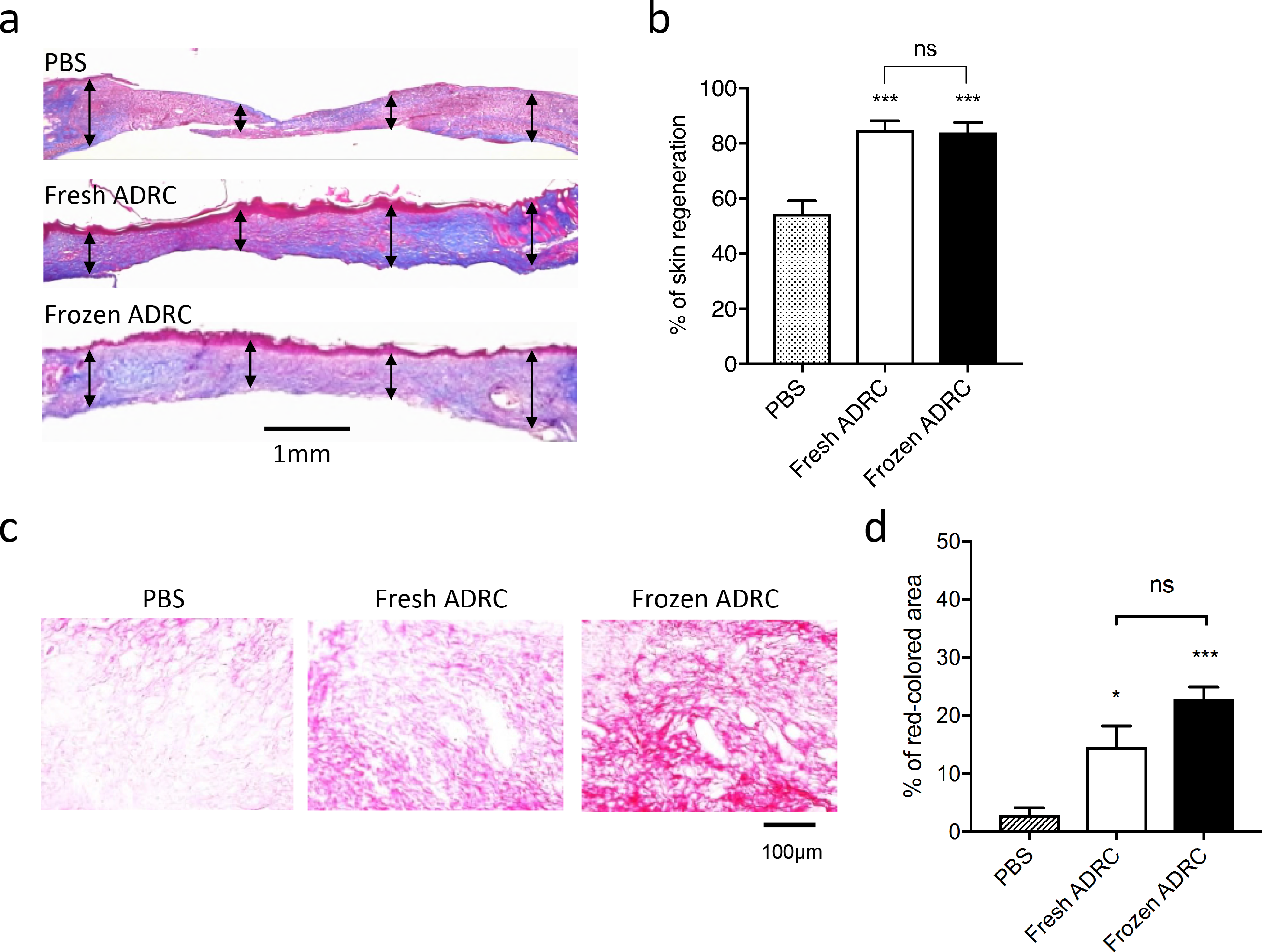
Histological analysis of the regenerated skin tissue. A, The tissue sections were stained with Masson’s Trichrome method on day 12 after surgery. Representative images of skin wounds are demonstrated. Double-arrow lines indicate skin thickness in each site. B, Quantitative analysis of tissue thickness. The ratio of regenerated tissue thickness to the original skin tissue thickness was calculated. C, The tissue samples were analyzed by Picro-Sirus red staining on day 12 after surgery. Representative images of skin wounds are demonstrated. D, Quantitative analysis of collagen production. The percent of the area of collagen fibers to the original area was calculated. ns; *, p < 0.05; vs. PBS group (control).

### Frozen ADRCs increased type I collagen production

The representative images showing collagen fibers under a standard light microscopy, (Figure 5C) and type I/III collagen under a polarized light microscopy. (Figure 6A) The percent of the area of collagen fibers to the original area was calculated. There was significant difference in the percent of collagen fibers between the ADRC groups and PBS group, but was no significant difference between fresh ADRC group and frozen ADRC group (Figure 5D). The percent of type I collagen fibers was significantly higher than that in fresh ADRC and PBS group (Figure 6B). In contrast, the percent of type Ⅲ collagen in PBS group, fresh ADRC group and frozen ADRC group (Figure 6C) demonstrated no significant differences. The percent of type I plus type III collagen fibers in frozen ADRC group was significantly higher than that in fresh ADRC group and PBS group (Figure 6D).

**Figure 6.**
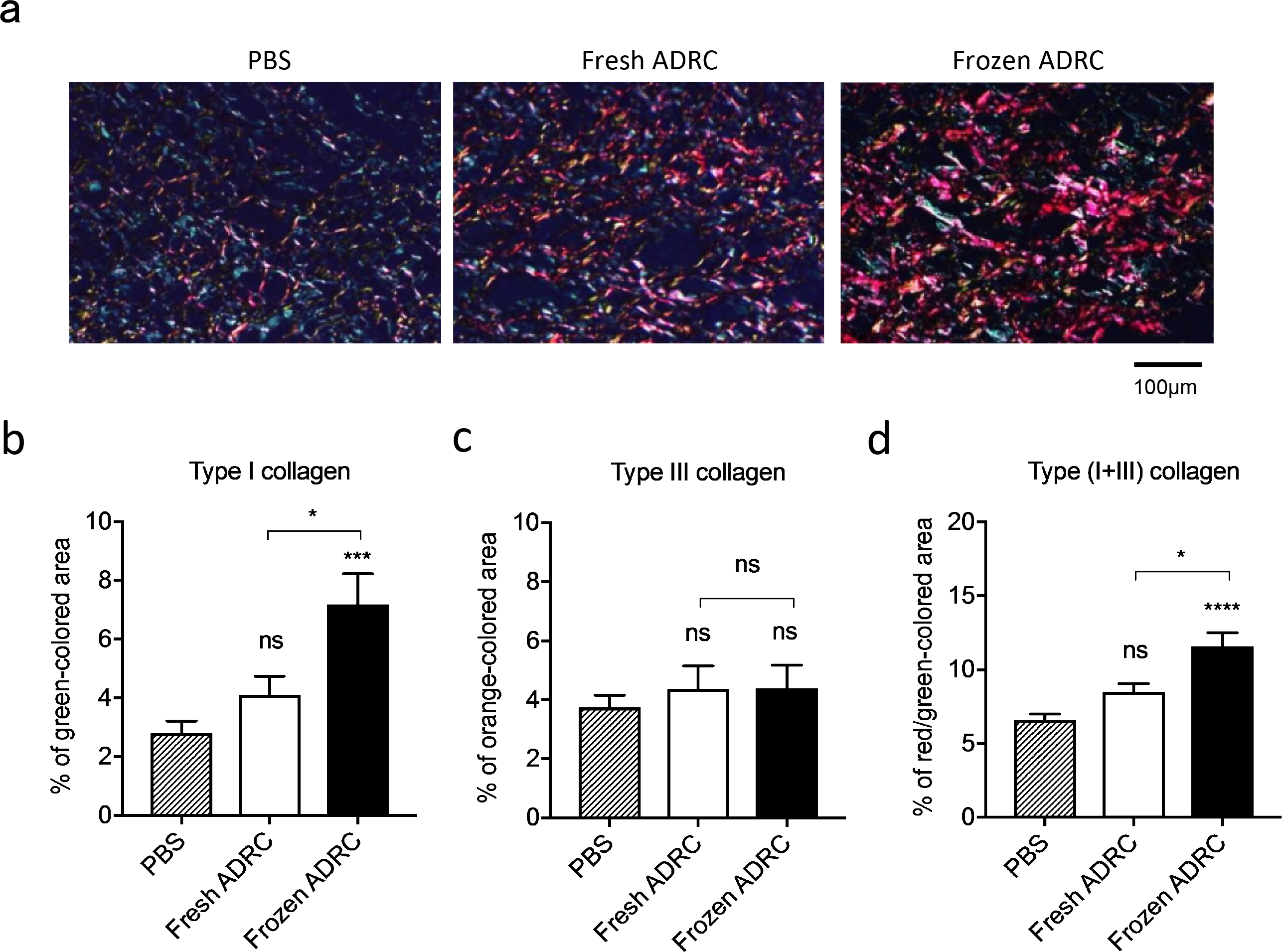
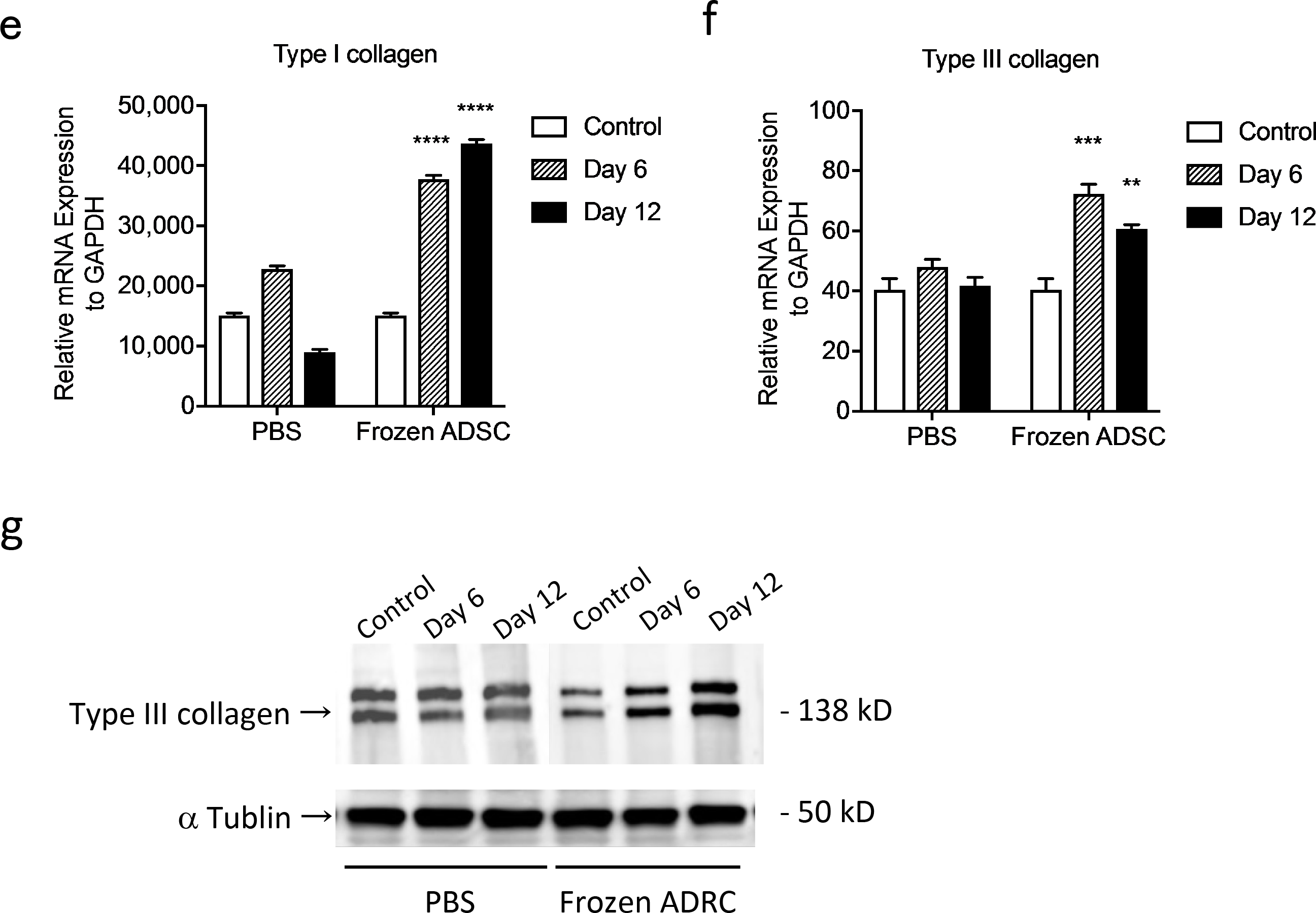
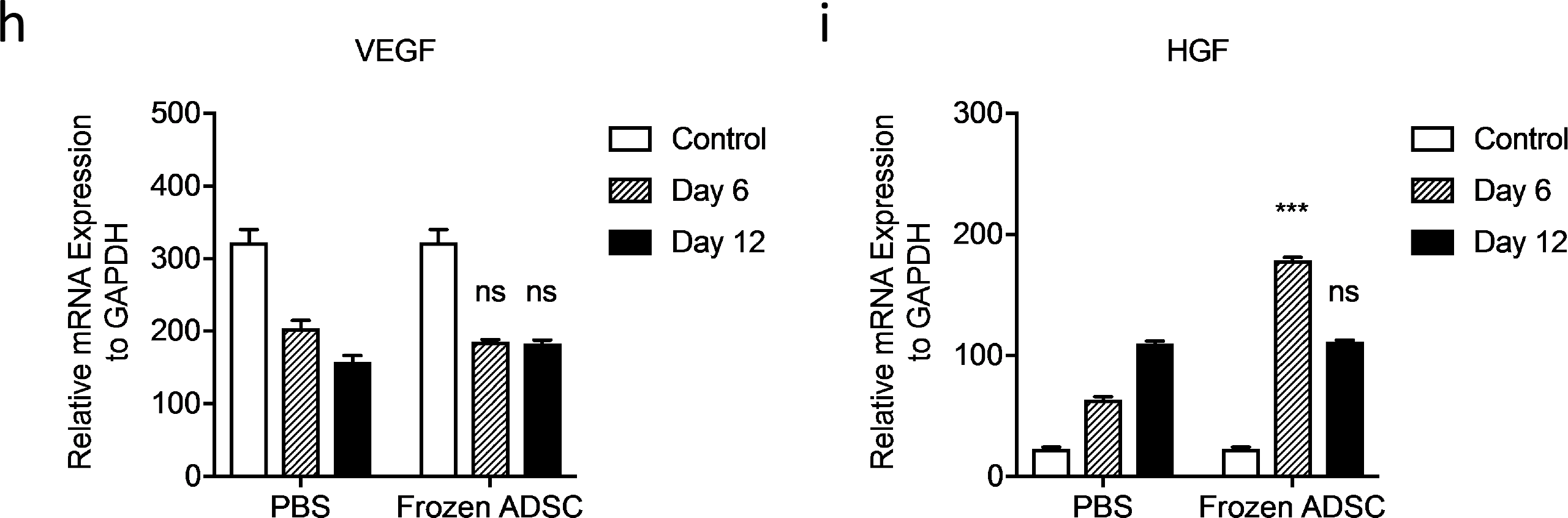
Assessment for collagen synthesis in regenerated wound tissue. A, The representative images of tissue section stained with Picro-Sirus Red in regenerated wound tissue under a polarized light microscopy. Type I collagen fibers are visualized in orange-red color and type III collagen fibers are visualized in blue-green. The percent of the area of each type of collagen fibers to the original area was calculated. B, Analysis of type I collagen fibers (orange-red). C, Analysis of typeⅢcollagen fibers (blue-green). D, Analysis of type I plus type III collagen fibers (orange-red+blue-green). The mRNA expression levels in regenerated skin tissue with either PBS or frozen ADRCs at day 6 and day 12 following surgery/treatment were examined by quantitative real-time RT-PCR analysis using specific primers for human genes of *type I collagen (COL1A1, E), type III collagen (COL3A1, F), VEGF (H) and HGF (I)*. The relative mRNA expression levels of target genes were normalized to the levels of *GAPDH* mRNA. ns; *, p<0.05; **, p < 0.01; ***, p<0.001; and ****, p < 0.0001 vs. PBS group (control). The production of type III collagen was further assessed by Western blot analysis. (G)

Next, we examined several gene expressions at day 6 and at day 12 both in PBS group and in frozen ADRC group. Although the gene expression level of type III collagen was overall low, those of type I collagen (COL1A1) and type III collagen (COL3A1) at both day 6 and day 12 were significantly higher in frozen ADRC group compared with PBS group. For pro-angiogenic growth factors, the gene expression of HGF was significantly high in frozen ADRC group compared with that in PBS group at day 6, but not at day 12. There was no significant difference of VEGF gene expression between PBS group and frozen ADRC group both on day 6 and day 12. (Figure 6E, F, H, and I) The gene expressions of FGF2 and EGF were not detected. Western blot analysis exhibited that the increase of type III collagen production was observed both at Day 6 and at Day 12 which was consistent with the results of gene expression as well as that of type I collagen. (Figure 6G)

### Frozen ADRCs increased neovascularization in regenerated skin tissue

Immunostaining for isolectinB4 (ILB4), a marker of endothelial cells, was performed to evaluate neovascularization. Angiogenesis was determined by directly counting the ILB4-positive dots. Representative images show ILB4-positive tissues at day 12 after treatment (Figure 7A). The direct counts of ILB4-positive dots had a significant difference between ADRC groups and PBS group. However, no significant differences was observed between fresh ADRC group and frozen ADRC group. (Figure 7B) There was a significant difference in direct counts of vessels greater than 5 μm in diameter between the ADRC groups and PBS group. No significant difference was observed between fresh ADRC group and frozen ADRC group. (Figure 7C) It was suggested that ADRCs induced angiogenesis in regenerated skin tissue and frozen ADRCs exerted similar effects on angiogenesis to fresh ADRCs.

**Figure 7.**
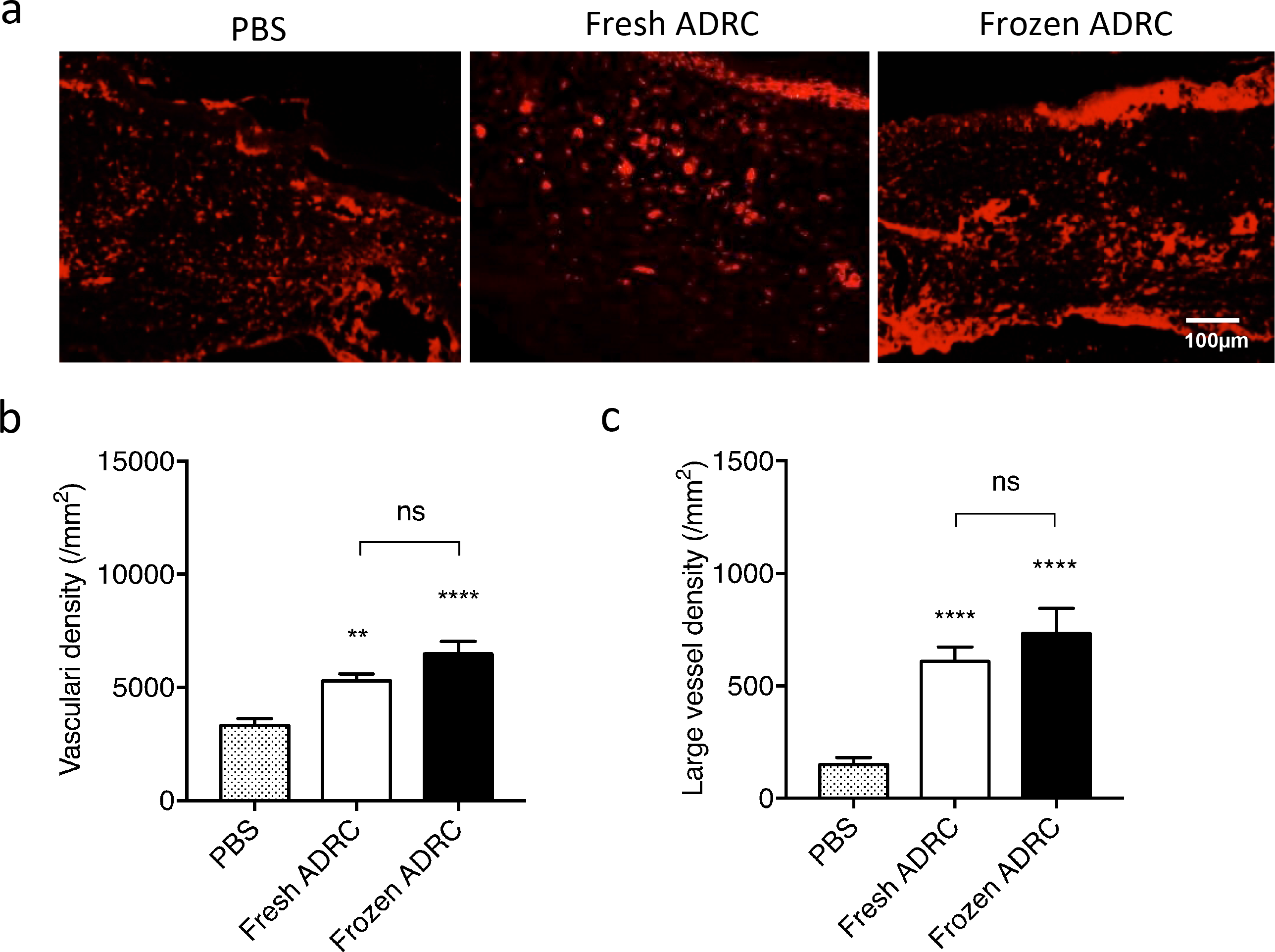
Assessment of vascularity in regenerated wound tissue. A, Fluorescent immunostaining for ILB4 (red) in the regenerated skin tissue. B, Analysis of entire blood vessels. All ILB4-positive dots and circles were counted. C, Analysis of large vessels. ILB4-positive large vessels (circles) with greater than 5 μm diameters were counted. ns; *, p < 0.05; **, p < 0.01; ***, p < 0.001; and ****, p < 0.0001 compared with the PBS group (control).

### Frozen ADRCs promoted tissue regeneration by paracrine factors

We examined the regenerated tissue including wound bed by fluorescent immunostaining with an anti-human mitochondria antibody to detect transplanted human cells. However, no transplanted cells were observed even on day 6 following application of cells with artificial dermis, suggesting that the endogenous tissue regeneration was promoted by the paracrine factors such as FGF2 which was highly expressed in frozen ADRCs (Figure 3B) but not by migration or direct contribution of the applied cells from the artificial dermis to regenerated tissue.

## Discussion

Although recent advances in burn care have significantly reduced overall mortality, new approaches in the treatment of burn wounds are still needed. In this regard, cell therapy has led to the development of innovative burn treatments [13]. The therapeutic effects of MSCs on burn wounds have been shown in a number of studies [13–16, 19–21]. Furthermore, the favorable effects of freshly isolated ADRCs, which are equivalent to SVF, on acute burns have been reported [15,16]. In contrast, the effects of cryopreserved ADRCs on burn wound healing have yet to be investigated thoroughly.

In this study, frozen ADRCs were hypothesized to improve burn wound healing to a similar extent as fresh ADRCs. Frozen ADRCs exerted similar therapeutic effects on burn wounds as fresh ADRCs in vitro and in vivo. We report the following main findings: 1) frozen ADRC-derived CM stimulated the proliferation and migration of keratinocytes and fibroblasts in vitro, and frozen ADRCs rather expressed high levels of growth factor genes, similar to fresh ADRCs; and 2) frozen ADRCs reduced the area of burn wounds by regenerating the dermal tissue and promoting vascularization to a similar extent as fresh ADRCs.

Keratinocytes and fibroblasts were cultured with CM derived from fresh and frozen ADRCs to identify the paracrine effects of fresh and frozen ADRCs on burn wound healing. The CM derived from ADRCs provided a migratory and proliferative stimulus for keratinocytes and fibroblasts. These results were consistent with previous studies [22,23].

The CM derived from ADRCs contains a fraction that is rich in soluble factors with paracrine actions, such as EGF, FGF, IGF-1, VEGF, and PDGF, cytokines important for wound healing [24]. The expression levels of the *FGF2*, *VEGF*, and *EGF* genes in ADRCs, particularly the *EGF* and *FGF2* genes, were significantly increased in frozen ADRCs compared with fresh ADRCs. Taken together, both fresh and frozen ADRCs appeared to be safe and feasible therapeutic tools without significant adverse effects on burn wounds. The important parameters for wound healing are dermal deposition and vascularization. ADRCs and ADRCs administration have been implicated in constructing the dermal tissue and neovascularization in burn wounds. We analyzed the dermal tissue thickness and counted ILB4-stained cells to evaluate dermal regeneration and neovascularization. The dermal tissue was thicker and many new vessels had developed in the fresh and frozen ADRC groups. Thus, ADRCs activated the proliferation and migration of fibroblasts and promoted angiogenesis. Based on the gross observation of the wounds, the efficacy of frozen ADRCs was the similar to fresh ADRCs. ADRCs also activated cell functions of fibroblasts promoting collagen synthesis regardless of freshly isolated or frozen stored cells. Specifically, type I collagen was significantly increased in frozen ADRC group, suggesting that frozen ADRCs promote wound healing with mature tissue [25]. The type I collagen production is positively regulated by FGF2 and EGF in fibroblasts, and EGF is also known to stimulate fibroblast replication with type I collagen accumulation during wound healing [26]. Our data of the EGF gene upregulation therefore accounts for the effect of frozen ADRCs on skin regeneration with mature tissue.

Thus, a trend toward better effects on the functions of keratinocytes and fibroblasts and angiogenesis was observed in the frozen ADRC group compared with the fresh ADRC group. The reason for the trend may be due to the potential purification of fresh ADRCs through cryopreservation and thawing, which eliminated miscellaneous cell populations other than MSCs, such as hematopoietic lineage cells. The flow cytometry analysis revealed the heterogeneity of fresh ADRCs, consistent with previously published studies [27]. Marker expression on frozen ADRCs changed, as shown by the increased frequency of CD90- and CD29-positive cells and the reduced frequency of CD34-positive cells. Based on our results, ADRCs were purified during cryopreservation and thawing, as the ratio of CD45+/11b+ cells decreased after thawing because neutrophils are sensitive to freezing and thawing [28].

In this study, frozen ADRCs exerted therapeutic effects on burn wound healing, although several studies have reported that fresh ADRCs accelerated burn wound healing.^15, 16^ Importantly, to our knowledge, this study is the first to show that frozen ADRCs improved burn wound healing to a similar extent as fresh ADRCs. Although the delivery methods for ADRCs deserve further investigation, these cells display the following advantages: 1) frozen ADRCs are convenient because the cells can be used as patients need them, 2) aliquoted cells can be used repeatedly within a certain interval depending on the time frame in which burns heal, and 3) thawed cells can be used very quickly without any isolation procedures.

## Conclusions

The treatment with frozen ADRCs resulted in sufficient therapeutic effects on burn wounds, including dermal synthesis and epithelialization along with increased vascularity, to a similar extent to fresh ADRCs. These findings may lead to the development of an innovative burn therapy using frozen aliquoted ADRCs in combination with conventional treatments, such as debridement and skin grafting.

## Acknowledgements

We thank Ms. Chinatsu Shiraoka for providing excellent technical assistance in the histological examinations, Mr. Yoshinori Koishi for providing excellent assistance in the animal experiments, and Sobajima Clinic for providing aspirated human fat tissue for this study.

